# Growth of the *Fucus* embryo: insights into wall-mediated cell expansion through mechanics and transcriptomics

**DOI:** 10.1101/2020.01.29.925107

**Authors:** Marina Linardić, Shawn J. Cokus, Matteo Pellegrini, Siobhan A. Braybrook

## Abstract

Morphogenesis in walled organisms represents a highly controlled process that involves cell proliferation and expansion; cell growth is regulated through changes in the structure and mechanics of the cells’ walls. Despite taking different evolutionary paths, land plants and some brown algae exhibit developmental and morphological similarities; however, the role of the algal cell wall in morphogenesis remains heavily underexplored. Cell expansion in plants is hypothesized to involve modifications of hemicellulose linkages and pectin gelation in the cell wall. Little is known about the wall-based control of cell expansion in brown algae; however, the algal analog to pectin, alginate, exhibits different gelation depending on its biochemistry. Here we show that cell wall mechanics and alginate biochemistry are correlated with cell expansion versus proliferation in the developing *Fucus serratus* embryo. In the elongating cells of the embryo rhizoid, we found a reduced cell wall stiffness and lower amounts of ‘stiffer’ alginate epitopes. In comparison, the early embryo thallus was shown to undergo cleavage-type cell proliferation, without expansion, and this was correlated with higher amounts of ‘stiff’ alginate epitopes and increased wall stiffness. An embryo development RNAseq dataset was generated to characterize differential gene expression during development. This data set allowed for identification of many enriched GO functions through developmental time. In addition, the transcriptome allowed for the identification of cell-wall related genes whose differential expression may underlie our observed growth phenotypes. We propose that differential gene expression of genes involved in alginate stiffness are strong candidates underlying differential wall stiffness and cell elongation in the developing *Fucus* embryo. Our results show that wall-driven cellular expansion mechanisms in brown algae are similar to those observed in plants. In addition, our data show that cleavage-type cell proliferation exists in brown algae similar to that seen in plant and animal systems indicating a possible conserved developmental phenomenon across the branches of multicellular life.

## Introduction

The formation of shapes has been of interest in the field of developmental biology for a long time, both in animal and plant systems (1). In walled multicellular organisms, such as plants and algae, morphogenesis results from cell expansion, division, and differentiation (2, 3). In plants, cell expansion is regulated mechanically by the balance between cell wall material and internal turgor pressure; when the balance tips, cell expansion results (4). A similar mechanical truth seems to exist for brown algal cells, given recent findings in the tip-growing cells of a model species, *Ectocarpus siliculosus* (5, 6).

In plants, cell expansion has recently been correlated with changes in wall matrix: the gel within which cellulose and hemicellulose fibers are embedded. In the model plant *Arabidopsis thaliana,* the biochemical and linked mechanical, properties of the pectin homogalacturonan within the cell wall matrix have been shown to regulate the magnitude of cell expansion (7–14). This may provide a direct mechanism for pectin rigidity to regulate cell expansion, but indirect effects from movement of other molecules within the cell wall (such as cellulose fibers or modifying proteins) may also be relevant (14, 15). Multicellular brown algal bodies are built from walled cells (16), whose cell walls are composed of cellulose, matrix polysaccharides, and proteins (more information on their biosynthesis and diversity can be found here (17, 18)). With cellulose content estimated at 1%-20% (19, 20), it is hypothesized that the wall gel matrix might be important for mechanical regulation of growth, similar to plant pectin (15,21,22).

Multicellular brown algae share a striking amount of morphogenetic similarity with plants and are often mistaken for plants. Similar developmental patterns such as from organ arrangement (23) and embryo patterning (24–28) to cell expansion types (e.g. tip growth (6), has led to this common misconception; however, brown algae are only very distantly related to the Viridiplantae (i.e. plants and green algae) and are estimated to have appeared ≈200 million years ago (29) through an independent evolutionary path. Since both plants and brown algae have cell walls with calcium cross-linked gel matrices, but independent evolutionary paths, the comparison of wall-regulated morphogenesis on a physical level may prove interesting; perhaps a similar set of physical rules regulating expansion might be operating. In the model plant *Arabidopsis thaliana*, cell expansion and division are critical for patterning the young embryo body after the first asymmetric cell division (30). Conversely, there is very little data on brown algal embryogenesis and its regulation.

Brown algal zygotes and embryos have served as a system to explore early morphogenetic events, such as polarity and asymmetric division since they fertilize and develop free from maternal tissue. The maternal-free aspect provides experimental access not found in plants, and *Fucus* has been used as a model species to study polarity and asymmetry during early embryo development (31–36). The initial asymmetric cell division produces two cells - a rhizoid and a thallus cell - with distinct morphologies and fates. The rhizoid cell generates the holdfast which will attach the alga to substrates, as well as the the stipe (stem) of the mature alga. The rest of the algal body (fronds, air bladders, and fertile structures in *Fucus*) develops from the thallus cell (25).

In *Fucus*, the cell wall matrix polysaccharides are sulfated fucans and alginate (17,18,37,38). It has been suggested that sulfated fucans play an important role in *Fucus* zygote polarization (39–41) but any role in cell expansion has yet to be uncovered. Currently, alginate seems a strong candidate for a matrix polysaccharide in brown algae that could regulate cell expansion. Alginate comprises the majority of the brown algal cell wall (**≈**50%; (19, 20). Like homogalacturonan in plants, alginate can cross-link with calcium, and thus regulate gel rigidity (21). Despite this mechanical similarity, alginate is distinct from pectin in structure as it is a linear polysaccharide composed of β-1,4-D-mannuronate (M) and α-1,4-L-guluronate (G) (42), produced as a mannuronate chain whose individual sugars can be epimerized to guluronate by mannuronan C-5 epimerases (43–45). It is the contiguous regions of guluronate, G-blocks, that form “egg-box” crosslinks with positively charged cations (such as Ca^2+^) leading to gelation (21, 46). The mechanical properties of the alginate gel, and therefore presumably the *Fucus* cell wall, are thus dependent on the amount of M and G sugars present: mixed MG regions are most flexible, followed by M-rich regions, with G-rich regions being the stiffest (MG flexibility > MM > GG; (47)).

Since *Fucus* affords a maternally-free developing embryo, it is an ideal system for studying the mechanics of morphogenesis in brown algae, and, specifically, that underlying cell expansion.

Here, we explore the mechanical basis of wall-mediated growth in the *Fucus serratus* embryo through a combination of atomic force microscopy and alginate immunohistochemistry. Furthermore, we present the first brown algal embryo development transcriptome and explore the expression of cell wall biosynthesis and modification genes in early embryo growth. We utilize our data to hypothesize that cell expansion in the *Fucus* zygote is regulated, in part, by alginate biochemistry and resulting wall mechanics. Our findings point to a physical similarity between the mechanical regulation of cell expansion in plants and brown algae.

## Results

### The *Fucus* embryo exhibits distinct growth behaviors between the rhizoid and thallus

The exploration of morphogenesis and growth in *Fucus* has focused mainly on the earliest events of polarization and asymmetric division (26,27,31,39,48–52). As such, no significant quantitative data on growth exists for embryo development beyond the first few days after fertilization (DAF). In order to explore *Fucus* embryo morphogenesis further, we examined embryo growth for 7 DAF at the organism and cell levels.

Using light microscopy, we first characterized the growth of the *Fucus* embryo on an organismal level. The embryo elongated over time at a decresing rate, reaching a length of ∼ 600 µm by 7 DAF (Fig. 1A) with initial rapid elongation leveling out between 3–7 DAF (Fig. S1A). Upon closer examination, embryos exhibited a highly consistent pattern of growth, distinct between the thallus and rhizoid body organs (T and R in Fig. 1A). Qualitatively, the thallus did not appear to elongate or increase in surface area for the first 5 DAF (Fig. 1A/B yellow); after 5 DAF, thallus began growing (Fig. 1A purple). In contrast, the rhizoid elongated from the beginning and contributed to the majority of to the embryos’ growth in length over the seven days (Fig. 1A/S1B green). Rhizoid elongation was directional with tip-like growth (Fig. S1C). As such, the leveling off (circa 3 DAF) of the whole organism elongation rate likely results from a decrease in rhizoid elongation, as observed qualitatively in our images (Fig. 1A green). After the drop in rhizoid elongation, embryos appeared to maintain elongation through the onset of thallus elongation (Fig. 1A/B purple). From these observations, we conclude that the *Fucus* embryo displays three distinct growth phases during 0-7 DAF: (i) an early phase dominated by rhizoid elongation, (ii) a middle phase where rhizoid elongation slows and thallus expansion initiates, and (iii) a late period dominated by thallus expansion.

**Figure 1.**
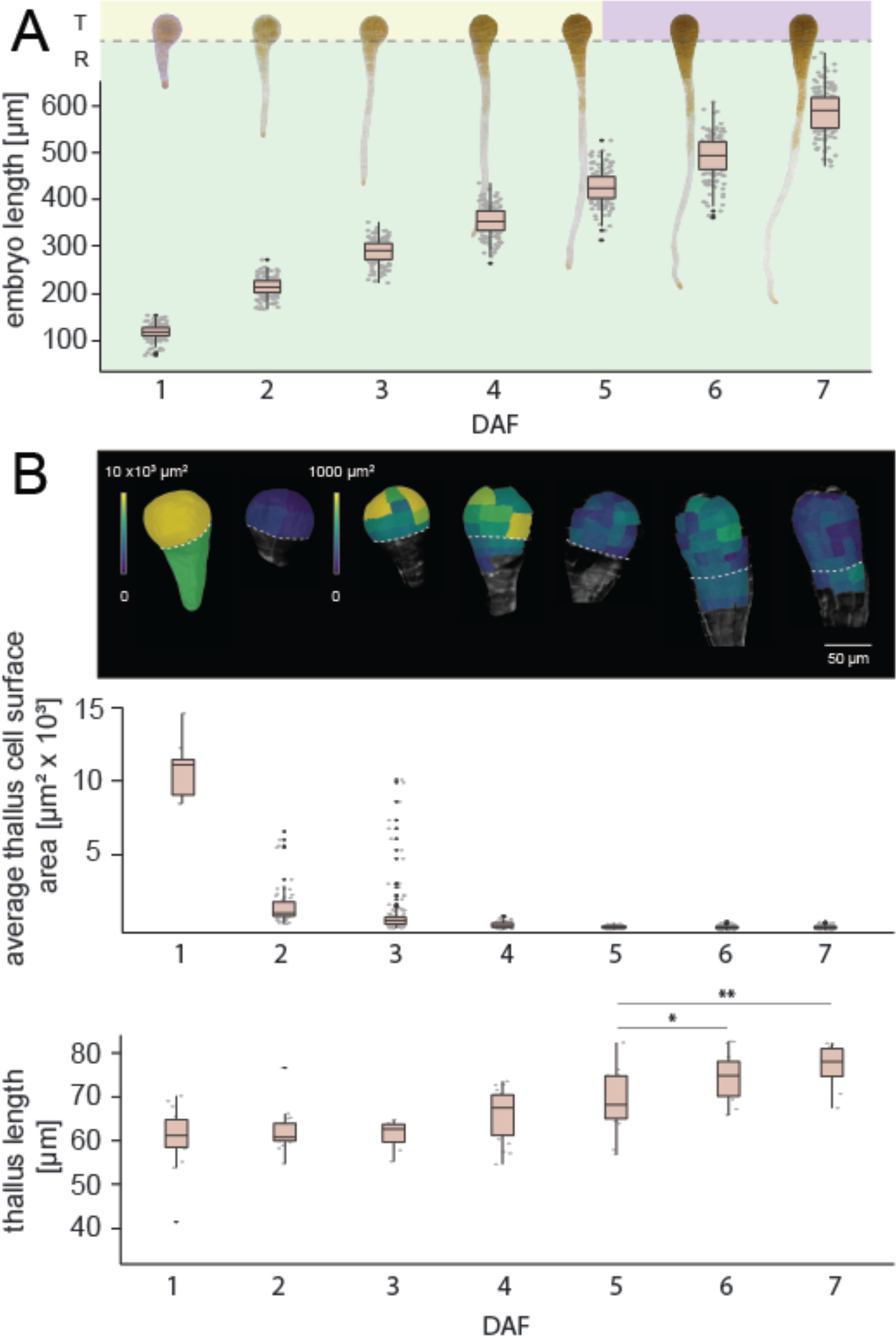
*Fucus* embryo growth dynamics. (A) Representative images of *F. serratus* embryo in the first seven days of development. T - thallus, R - rhizoid. Graph shows the embryo length increase in time. Embryo growth can be divided in three phases: rhizoid elongation (green), exclusive thallus division only (yellow) and thallus expansion (purple). Embryos observed, N=150. (B) Temporal quantification of average surface areas of single cells derived from the initial thallus cell. Representative heat maps of cell surface areas are depicted first, with graphical quantification below. Embryos observed, middle panel: N(1DAF)=9; cells measured=9, N(2DAF)=9; cells measured=62, N(3DAF)=11; cells measured=151, N(4DAF)=9; cells measured=119, N(5DAF)=8; cells measured=194, N(6DAF)=9; cells measured=274, N(7DAF)=8; cells measured=274, embryos observed, bottom panel: N(1DAF) =15, N(2DAF) =17, N(3DAF) =7, N(4DAF) =21, N(5DAF) =15, N(6DAF) =16, N(7DAF) =10. Significance at *p<0.05, **p<0.01 according to the pairwise Student’s t-test (normal distribution, equal variance).

To examine the contribution of cell-level growth (i.e., division and expansion) to the organ-level growth patterns we observed, we quantified cell shape and size over time. For this, we followed cell wall staining with confocal imaging and MorphoGraphX analysis (53). In our hands, staining with calcofluor white yielded our best cell outlines in developing embryos, allowing visualization and quantification of cell surface areas. However, we were only able to image cell surfaces in epidermis, despite numerous methodological attempts, including fixation and clearing as employed by Yoshida et al. (30) in the Arabidopsis embryo. At 1 DAF, the expected asymmetry in thallus and rhizoid initial cells was evident (Fig. 1B). We therefore proceeded to time sample embryos and analyze epidermal cell division and expansion in a pseudo-growth series.

While our light microscopy had indicated little expansion in the thallus up to 5 DAF, we were able to see cell divisions occurring in this tissue as early as 2 DAF (Fig. 1B). Thallus divisions occurred rapidly on the embryo surface (Fig. 1B). These divisions, however, did not seem to increase the thallus size before 5 DAF (Fig. 1B) and instead resulted in a decrease in the surface area of daughter cells with a plateau achieved circa 5 DAF (Fig. 1B).

This suggests that, in the early stages of embryo development the thallus undergoes cleavage-type divisions, with little to no cell expansion. This phenomenon is rare in plant development but observed in the early embryos of the plant *Arabidopsis* and in metazoan embryos (30, 54). After 5 DAF, cell surface area expansion was observed coincident with cell division yielding a near-constant mean cell surface area (5 – 7 DAF cell surface area = 242 ± 22 µm^2^; Fig. 1B). Note that 5 DAF mark is also where our light microscopy indicated that the thallus began elongation (Fig. 1A). Note that 5 DAF corresponds to visible appearance of apical hairs (Fig. S1D); apical hair development is linked to meristematic cell establishment and is necessary for further embryo growth (55). As such, it seems likely that the establishment of a meristematic cell in *Fucus* is necessary for volumetric thallus growth. We thus conclude that the *Fucus* thallus initially undergoes cleavage-type divisions but soon transitions to a combination of cell expansion and division that yields a growing organ.

Confocal observations of growing rhizoids indicated that rhizoid cells underwent expansion from 1 DAF, followed by divisions (Fig. S1B). Initial single rhizoid cell grew by tip-growth (Fig. S1C), and as the rhizoid extended, several perpendicular divisions occurred. Tip-growth has recently been described for the filamentous brown alga *Ectocarpus* (6). Interestingly, the filamentous cell in *Ectocarpus* may exhibit tip-growth due to a thinning of the cell wall near the tip as opposed to the material composition differences as in plants and fungi (6); as a brown alga, *Fucus* may share a similar tip-growing mechanism to the one in *Ectocarpus*. The sum rhizoid cell surface area showed a rapid increase within the first 2 DAF (Fig. S1B) with a slowing over time terminating with holdfast production (∼10 DAF, data not shown); recall that our organ-level elongation rate began decline by 3 DAF (Fig. 1A). These data are consistent with an early and rapid rhizoid elongation supported by cell division.

Taken together, our data paint the following picture of *Fucus* embryo growth over the first 7 DAF: after asymmetric division at 1 DAF, the rhizoid elongates rapidly through expansion and division for the first 3 DAF. During this time, the thallus undergoes cleavage-type divisions but does not increase appreciably in organ size. At 3 DAF, rhizoid elongation begins to slow, likely in preparation for holdfast formation. During this interval, it is likely that the meristematic apical cell was established within the thallus, leading to the growth of the thallus starting at 5 DAF via cell expansion and division. As such, there are three growth phases evident in the early *Fucus* embryo – (i) rapid elongation and division in the rhizoid (ii) cleavage divisions in the thallus, and (iii) coupled elongation and division coupled within the thallus. The combination of these growth modes yields the characteristic *Fucus* embryo morphogenesis.

### Embryo growth phases correspond to changes in tissue mechanics

In plants, cell expansion has been correlated with changes in cell wall mechanics that allow the cell wall to yield to turgor pressure, resulting in growth (15,56,57). Given the presence of cell walls in the *Fucus* embryo, we wondered whether cell wall mechanics might underlie the different growth modes observed, specifically cleavage-type volumetric division in the thallus versus expansion followed by division in the rhizoid. To assess cell wall stiffness in developing *Fucus* embryos, we employed Atomic Force Microscopy (AFM) based indentation. AFM indentation examines the force required to deform cell wall material within a given area providing spatial resolution. This technique was applied to embryos at 3 DAF when the thallus and rhizoid displayed differential growth behavior; indentations were performed along the longitudinal axis of the embryos so as to obtain information about both thallus and rhizoid from each sample.

Analysis revealed that at 3 DAF the *Fucus* thallus was stiffer than the rhizoid (Fig. 2A/B). This observation correlates wall stiffness with cell elongation, providing support for a relationship similar to that seen in plants: stiffer cell walls reduce cell elongation. We performed additional AFM-based stiffness measurements at 1 and 10 DAF (Fig. S2) and observed that the rhizoid was consistently less stiff than the thallus. The thallus was at its stiffest in the 3DAF samples (Fig. 2A/B, S2) coincident with the cleavage-type divisions observed in the thallus at this stage of growth.

**Figure 2.**
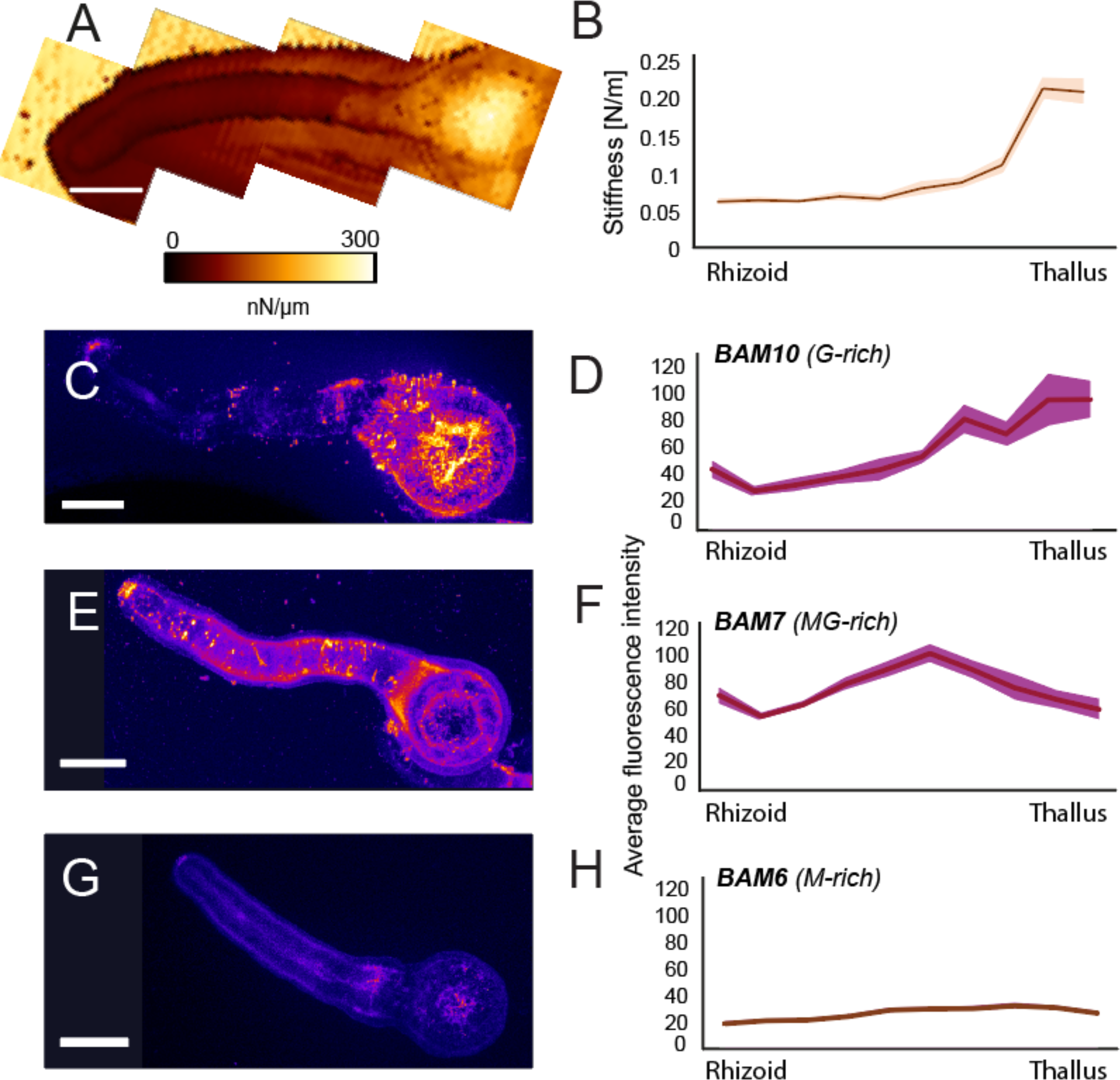
The *F. serratus* embryo displays stiffer cell walls with more G-rich alginate in the dividing thallus at 3 DAF. (A, B) Embryo stiffness displayed as a representative map (A) and quantitatively over all samples (n=7) along the embryo length (B). (C-H) *In muro* immunolocalization of alginate epitopes in 3 DAF embryos. Fluorescence indicates localization of BAM10 (C,D), BAM7 (E,F) and BAM6 (G,H) binding. (C,E,G) representative confocal images of immunolocalizations. (D, F, H) Fluorescence quantification from immunolocalizations along the embryo length, for all samples analyzed (N(BAM10) = 9, N(BAM7) = 12, N(BAM6) = 21).

### Alginate biochemistry links to wall mechanics and the *Fucus* embryo expansion pattern

Given the relationship between pectin biochemistry and wall stiffness in plants, and the biochemical relationship between alginate epimerization and gel stiffness (21), we next examined alginate biochemistry using monoclonal antibodies (39). Antibodies recognized guluronic (G)-rich areas (BAM10), mannuronic (M)-rich areas (BAM6), and mixed mannuronic-guluronic (MG) regions (BAM7). These types of alginate likely correspond to stiff, intermediate, and least stiff (58) alginate mechanical behavior. Whole-mount immunolocalizations were performed on fixed 3 DAF embryos when the difference between the thallus and rhizoid was highest in terms of growth behavior and wall stiffness. All three antibodies successfully reacted with the embryos and negative controls are shown in Fig. S3.

The epitope bound by BAM10 (G-rich areas) was detected more in the thallus than the rhizoid (Fig. 2C/D), indicating this region may contain stiffer alginate compared to the rhizoid. Conversely, the epitope of a softer alginate (BAM7; MG-rich areas, Fig. 2E/F) was detected at higher levels in the rhizoid and the rhizoid tip, compared to the thallus. Both BAM10 and BAM7 showed fluorescence in the rhizoid tip, as has been observed previously (5) and may relate to the secretion of the adherent matrix. BAM6 (M-rich areas, Fig. 2G/H) did not label the embryo strongly and its epitope was detected uniformly over the embryo. Our alginate immunolocalizations are therefore consistent with our stiffness observations in the embryo: the stiffer thallus presented more G-rich alginate while the less stiff rhizoid presented more of the less-stiff MG-rich alginate. These data support a model in the *Fucus* embryo where alginate biochemistry influences cell wall stiffness, which in turn influences cell expansion and overall embryo growth behavior.

### Transcriptional changes during Fucus embryo development: a transcriptomic approach

There are currently no means for transformation or mutagenesis in *Fucus* which would permit direct attack on the correlation between wall stiffness and growth. As such, we initiated a *de novo* transcriptomics approach to examine changes in gene expression during embryo development. We generated an embryonic transcriptome for *Fucus* from pooling three biological replicates for each of four stages of development: “*a*”, 7 hours after fertilization (7 hours after fertilization (HAF): round, fertilized, and a zygote without fixed polarity); “*b*”, 1 DAF (post-germination and first asymmetric division); “*c*”, 3 DAF (thallus divisions and rhizoid elongation); and “*d*”, 10 DAF (thallus growth and rhizoid holdfast formation). Thus, there were a total of 12 RNA-Seq samples.

Trinity assembly yielded 127,489 putative transcript isoforms/fragments which were, after processing (see Methods), considered to represent 24,691 protein-coding genes (albeit with some genes likely duplicated, fragmented, or partial, or with incorrect CDS vs. UTR partitioning/codon phasing or intron rejection). Amino acid alignments to proteins in NCBI found 67% (17K) as known (E-value < 10^−5^), with 76% of the knowns having a best hit to *Ectocarpus siliculosus*. The known count is similar to the 16K predicted genes of *Ectocarpus*, 14K of *Cladosiphon okamuranus*, and 19K of *Saccharina japonica* (59–61). 57% of best alignments involved at least half the sequence of both the *Fucus* and NCBI protein, and the median amino acid identity of *Fucus*–*Ectocarpus* alignments over *Fucus* genes with a best hit to *Ectocarpus* was 61%. (Thus, although *Ectocarpus* is the closest organism with good representation in NCBI, it is not that particularly close to *Fucus* in an absolute sense.) Expression of many genes changed drastically over the developmental timecourse; of the known genes, a null hypothesis of constant expression over time was rejected for 20% and 46% at *q*-levels 10^−5^ and 10^−4^, respectively.

To characterize differentially expressed genes, Gene Ontology (GO) enrichment analyses were performed, combining *Fucus* gene–to–GO term assignments from a run of InterProScan directly on our *Fucus* protein-coding genes with those transferred from *Ectocarpus* via orthologs as determined by a run of OMA on *Fucus*, *Ectocarpus*, and four other related organisms. For each of the 74 non-constant ways (“patterns”) the expression levels of the four timepoints (*a* = 7 HAF, *b* = 1 DAF, *c* = 3 DAF, *d* = 10 DAF) could be weakly ordered (i.e., *a < b < c < d* vs. *a = b > c = d* vs. … except *a = b = c = d*), hypergeometric GO enrichment p-values were determined for the subset of genes that could be statistically determined to be of that ordering (see Methods). 25 patterns had at least one GO term with p-value below 10– 4 (see Fig. S7/S8), with highlights summarized in Fig. 3. These patterns fall into six rough groups based on expression levels peaking at a particular timepoint (*a, b, c,* or *d*) or dipping at a particular timepoint (*b* or *d*).

**Figure 3.**
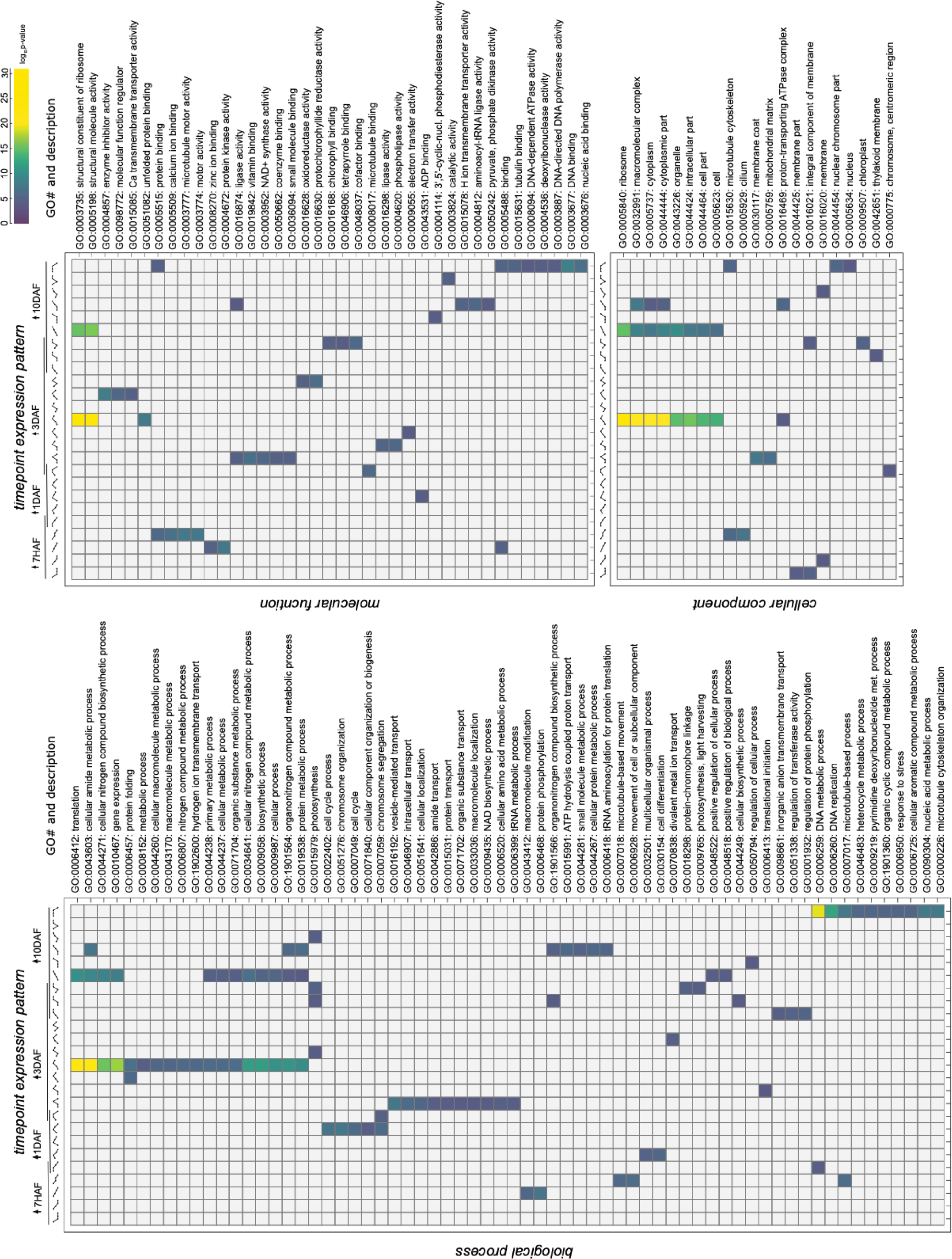
Highlights of Gene Ontology (GO) enrichments in different temporal expression patterns. Represented here are terms separated by the GO root term category: Biological Process, Molecular Function, and Cellular Component. X-axis represents the pattern of developmental expression indicated by spark-lines; 4 dots connected with a line that changes according to the increase or decrease of expression with respect to the other time points. Every illustration follows the same rule: first dot 7 HAF, second dot 1 DAF, third dot 3 DAF, and fourth dot 10 DAF; enrichment at a specific timepoint is shaded in grey and the overlaps represent enrichment in both neighboring time points. The *p*-value for GO terms ≤0.0001 is illustrated in the heat map. For full heat map including all rows and columns and all terms with *p*-value ≤ 0.0001, see Fig. S7.

Patterns peaking at 7 HAF included terms related to microtubule-based movement, protein phosphorylation, ion binding, and membrane. Note that as the zygote is getting ready to asymmetrically divide, it undergoes polar distribution of cellular components, engaging the cytoskeleton (26,34,62,63). Enriched terms specific to 1DAF were related to ‘cell cycle’, ‘chromosome organization’ and ‘cell differentiation’. These enrichments also make sense in terms of embryo biology since the thallus and rhizoid fates are established here by asymmetric cell division. For peaking at 1 DAF and transitioning to peaking at 3 DAF, we find terms involving the cell cycle and cell differentiation, including chromosomes, centromeres, and their organization and segregation. These are processes one would expect to be enriched, continuing from 7HAF, as the early fucoid embryo undergoes cell divisions that require activation of mitotic cell cycle genes and cytoskeleton to give rise to two differentiated cells, thallus and rhizoid progenitors (24, 51).

Terms enriched in patterns peaking at 3 DAF or 10 DAF or highest at both included ones related to protein production/maturation, localization, protons/energy, binding of vitamins/coenzymes/cofactors, and photosynthesis and its machinery (including its construction). Indeed, as the embryo matures, the photosynthetic apparatus becomes more active and photosynthetic efficiency increases (64–66).

Peak expression patterns at 10DAF exhibited GO enrichment in terms similar to 3 DAF-peak ones such as ‘translation’, ‘metabolic process’, ‘DNA replication’, ‘gene expression’. Again, these GO enrichments are consistent with the ongoing maturation and development of the *Fucus* embryo at 10DAF.

Altogether, our embryo transcriptome is consistent with our knowledge of embryo developmental biology in *Fucus*. We expect that this resource will be valuable to the community as it represents the first developmental embryo transcriptome of a brown alga. Further analysis will prove crucial to understanding the whole of *Fucus* embryo development.

### Expression pattern of cell wall-related genes

As our previous observations suggested that the differential growth displayed between the thallus and rhizoid was correlated with cell wall stiffness and alginate biochemistry, we examined the timecourse expression patterns of genes related to cellulose, sulfated fucans, and alginate, as these play roles in cell wall biosynthesis and modification. A table of cell wall biosynthesis genes and their closest homolog in *Ectocarpus* may be found in Table S1.

Twelve genes in our *de novo* transcriptome had homology to cellulose synthases (Table S1; *Ectocarpus* CESA and CSL; (38)). All were differentially expressed in at least one time-point (Fig. 4A). Expression patterns varied, although most (like many of our genes) exhibited broad patterns of either general increase or general decrease over time (Fig. 4A). This suggests variable cellulose biosynthesis dependent on developmental stage.

**Figure 4.**
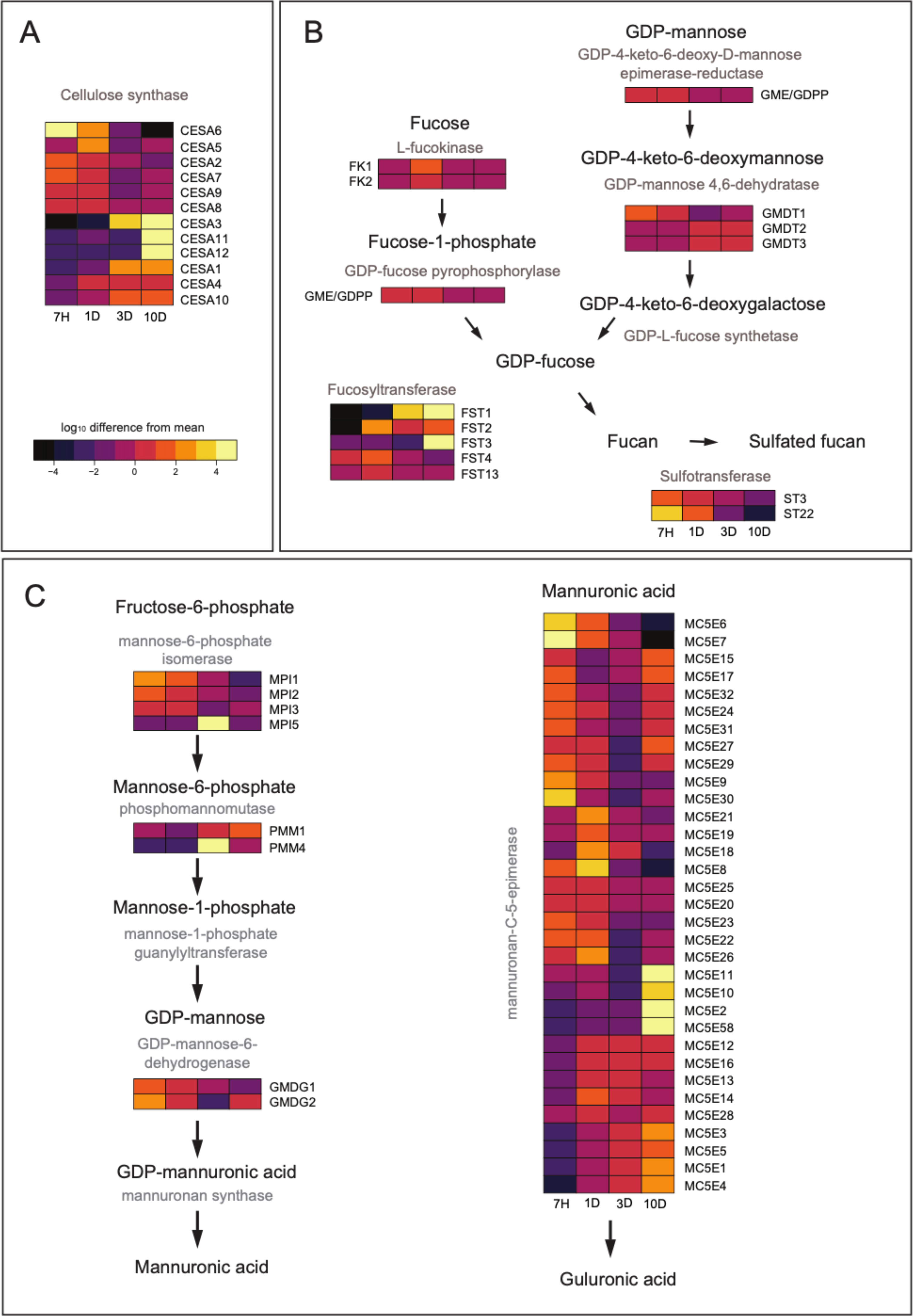
Schematic representation of the biosynthetic pathways of three main components of the brown algal cell wall: (A) cellulose, (B) sulfated fucans and (C) alginate, with expression levels of identified genes during *Fucus* embryo development (1 hour (H), 1 day (D), 3 days and 10 days after fertilization). Identifiers of putative relevant genes from the *de novo* transcriptome are shown, along with their relative expression levels across the *Fucus* embryo development timecourse; FK – fukokinase, GME/GDPP - GDP-4-keto-6-deoxy-D-mannose epimerase-reductase/GDP-fucose pyrophophorylase, GMDT - GDP-mannose 4,6-dehydratase, FST – fucosyltransferase, MPI – mannose-6-phosphate isomerase, PMM – phosphomannomutases, GMDG - GDP-mannose 6-dehydrogenase, MC5E – mannuronan C5-epimerase, ST – sulfotransferase, CESA – cellulose synthase (corresponding Trinity gene names found in Table S1.). Represented here are the genes with the differential expression in at least a one time-point (p-value<=0.0001). For the full set of putative homologs see Table S1.

The full biosynthetic pathways for sulfated fucans and alginate in brown algae are still uncharacterized, although there is progress (38). Following these proposed pathways, we found several genes (discussed below) putatively corresponding to enzymes in both sulphated fucan and alginate biosynthesis.

For sulfated fucans, a total of 56 putative homologs for the following enzymatic genes were identified: two L-fucokinases, one GDP fucose pyrophosphorylase, three GDP-mannose 4, 6-dehydratases, one GDP-4-keto-6-deoxy-D-mannose epimerase-reductase, four GDP-mannose 4,6-dehydratases, thirteen fucosyltransferases, and thirty-two sulfotransferases (five of which were homologous to carbohydrate sulfotransferases, Table S1.). Of these, only 14 were statistically inconsistent with constant expression over time (*q*-value < 10^−4^) and appear in Fig. 4B; full gene list in Table S1.). When examining the DE genes we could conclude the following: enzymes leading to the production of GDP-fucose showed both slight upregulation and downregulation during embryo development. Fucosyltransferase expression pattern varied depending on the gene, with three genes showing up-regulation in the later developmental stages. Both carbohydrate sulfotransferases showed a higher expression in the early stages (7H and 1DAF) which may be related to the cues of sulfated fucans required for both zygote polarity and adhesion (39–41).

Interpretation of gene expression in terms of the sole production of sulphated fucans is challenging to achieve here; most pathways have not been characterized in brown algae. Even though most biosynthetic pathways are not yet characterized in brown algae, GDP-fucose might indeed be a component of multiple metabolic pathways. In plants, GDP-fucose is found as a component of glycan structures such as N- and O-linked glycans, xyloglucans, pectins and arabinogalactan proteins (AGPs) (67–71). Some of these structures have been also found in brown algae, such as AGPs (72). In addition, transcript levels may not reflect levels of protein products (and enzyme activity), and such a difference has been previously reported to occur in the brown alga *Saccharina japonica* (73).

From the alginate biosynthesis pathway, a total of 14 putative homologs were found: five putative mannose-6-phosphate isomerases, four phosphomannomutases, and five GDP-mannose-6-dehydrogenases (Table S1.). As with the sulphated fucan biosynthetic homologs, only some varied in expression, and these appear in the left half of Fig. 4C. Isomerases and GDP-mannose-6-dehydrogenases showed higher expression in early embryo development, whereas phosphomannomutases had higher expression levels at 3 and 10 DAF (Fig. 4C). Alginate pathway products may also be used in other biosynthetic pathways; for instance, GDP-mannose can be a substrate for N-linked glycans in plants (74), as well as a substrate for sulfated fucan biosynthesis in brown algae; again, interpretation of expression is not completely straightforward.

The modification of alginate to alter stiffness, and the enzymes responsible, is most relevant to our data presented thus far. Once alginate is produced, it can be epimerized by mannuronan C5-epimerases (MC5Es), leading to changes in alginate gelling (75). We identified 59 putative MC5Es in our transcriptome (Table S1) of which 35 had non-constant expression (right half of Fig. 4C). These showed variability in expression patterns, but several genes displayed high expression early in embryo development and several had later peaks (Fig. 4C). To date, a number of bacterial genes encoding MC5Es have been discovered and the exact functions of some of these enzymes (their patterns of epimerization) have been investigated (76, 77). It is likely that each epimerase found here has a specific epimerization pattern and is required in different developmental stages, contributing to differential cell and tissue expansion. Further analysis of these newly identified putative epimerases will prove essential in understanding their role in *Fucus* embryo development, wall stiffness, and putative influence on cell growth behavior.

## Discussion

### An interplay between *Fucus* embryo wall mechanics and biochemistry on a cellular level

During embryogenesis, fucoid embryos have to coordinate a series of important developmental steps in order to ultimately create their adult body plan. Starting with polar axis formation and the first asymmetrical cell division, the division between the thallus and rhizoid is set. Manipulating these two events can heavily influence the growth of young embryos (50,78,79). The different growth phases we observed in the embryo provided an excellent model to examine the basis of organ and cell growth in the brown algal lineage, a highly diverse group of organisms that have only barely been explored.

Our data correlate wall mechanics and alginate biochemistry to cell expansion in the *Fucus* embryo. We report that at 3 DAF, the *Fucus* embryo displays differential growth behavior between the thallus and the rhizoid: the thallus is not expanding but is undergoing cleavage-type cell divisions while the rhizoid is rapidly elongating. This differential cell expansion behavior correlates with cell wall stiffness: the expanding rhizoid has a less stiff cell wall. Finally, both expansion and stiffness were correlated with alginate biochemical epitopes: less expanding cells, with stiffer walls, had more epitope signal from G-rich alginate epitopes; expanding rhizoid cells were less stiff and had higher MG-rich epitope signal for alginate. These data show that cell expansion in *Fucus* embryos is likely limited by cell wall mechanics, underlain in part by alginate biochemical modification by mannuronan C5-epimerases.

### The cell wall during *Fucus* embryo development

The cell walls of 24h old *Fucus* embryos consist of ∼ 60% alginate, 20% fucans, and 20% cellulose (20). Cellulose is the load-bearing polysaccharide in plants and algae and it has been reported that the arrangement of the cellulose microfibrils is correlated with the direction of the cell expansion in both lineages (80–84). In the *Fucus* zygote, cellulose has been suggested to act as a strengthening component as its low signal detected during germination corresponds to the growth initiation (85). Even though it may be involved in regulating anisotropic expansion, cellulose may not be a main contributor to the mechanical properties observed here, which has also been hypothesized for *Ectocarpus* (86).

In this study we have identified 12 putative cellulose synthase homologs expressed during embryo development in *Fucus*; further exploration of these genes, their products, and their roles in cellulose synthesis will be crucial to understanding how cellulose contributes to algal development. It is possible that different cellulose synthases are associated with distinct developmental processes, as has been shown in other walled organisms such as *Physcomitrella*, *Brachypodium*, and maize (87–89). In addition, we have identified both CESA and CESA-like homologs; CESA-like proteins in plants are involved in callose biosynthesis. Recent work indicates that β-1,3-glucans appear to be present in *Fucus* cell walls and they may be involved in callose biosynthesis (90). We must note that the major limitation here is the lack of molecular genetic tools in brown algae for forward genetic analyses of gene-phenotype relationships.

Sulfated fucans have been linked to zygote adhesion and stress tolerance (41, 52), but have also been suggested to strengthen the tip-growing rhizoid during elongation (91). In our data set, we identified several possible homologs involved in sulfated fucan biosynthesis whose exact roles deserve further exploration. In addition, we identified two putative carbohydrate sulfotransferases which are strong candidates for the final addition of sulfate to cell wall fucans. Exploration into these genes would be essential for understanding the role of sulfated fucans in embryo development and beyond.

The biosynthesis pathway of sulfated fucans did not show a clear pattern of up- or down-regulation during embryo development. We did, however, find an interesting observation; the *FucSerDN35869c0g1i2* gene was found to be highly homologous to both GDP fucose pyrophosphorylase (AUN86413.1, 93.6% identity) and GDP-4-keto-6-deoxy-D-mannose epimerase-reductase (Ec-01_003130.1, 94% identity). This observation could relate to the dual function of a single enzyme in the fucan pathway, although this is hard to confirm with only the transcriptome data generated here. There have been reports of other enzymes in the wall biosynthesis that are thought to perform a dual activity such as mannose-6-phosphate isomerase which could perform the function of mannose-1-phosphate guanylyltransferase (92). The amount of currently available molecular data on brown algal species is very scarce. These data further emphasize the need to continue exploring the brown algal lineage to be able to discern these ‘dual’ homologies and gain a better understanding of algal metabolism and their biosynthesis pathways.

Alginate has previously been reported to have an important mechanical role and has the ability to change the mechanical properties of materials depending on the formation of calcium bridges between guluronic acid-rich areas (21). The data here suggest that the G-rich alginate (as detected by BAM10) is more prevalent in the non-expanding thallus cells, whereas the ‘softer’ MG-rich alginate (as detected by BAM7) is found more in the actively expanding rhizoid area. Similar to the results from our study, a lower abundance of G-rich alginate was detected in the actively expanding cells of *Adenocystis utricularis* (93).

The alginate biosynthesis pathway has not yet been fully described in the brown algal lineage. However, a few genes homologous to bacterial genes involved in this pathway have been identified in *Ectocarpus and Saccharina* (38,61,92). In the *Fucus serratus* transcriptome, some of the genes seem to have a few more homologs. This might be due to the fact that our transcriptome is fragmented and they are actually a single gene, or there are multiple homologs that have the same function in *Fucus*. Furthermore, in the *Ectocarpus* genome, there was no homolog for mannose-1-phosphate guanylyltransferase (MPG) enzyme which catalyzes the reaction from mannose-1-phosphate to GDP-mannose. We have found a single homolog of the MPG in the *Fucus* transcriptome, matching with the *Arabidopsis* mannose-1-phosphate guanylyltransferase 1 (CYT1; 51%). It has been recently reported in *Saccharina japonica* that another enzyme has the ability to act as an MPG (92). However, the enzyme they report is homologous to a *Fucus* Trinity gene in our study (FucSerDN37605c0g2i1) which is different from our newly identified putative MPG (FucSerDN28326c0g1i1). This enzyme was not sufficiently expressed to be considered in the downstream gene expression analysis. However, we are currently working on generating a *Fucus* genome; this will enable us to look into more detail at these ambiguous genes that have not been identified in other brown algae, but have been identified in our analysis.

Here we found 62 homologs of mannuronan C5-epimerases with several expression patterns during development. Previous molecular analyses have revealed candidates for 31 genes in *Ectocarpus siliculosus* (Michel et al., 2010), 105 in *Saccharina japonica* (61), 6 in *Laminaria digitata* (94), and 31 in *Undaria pinnatifida* (95). The variety of potential epimerases found in the species of the brown algal lineage suggests that brown algae might have evolved the ability to ‘tweak’ the alginate structure to finer detail than what is observed in bacteria. The variability in the expression pattern in our dataset might reflect this hypothesis. However, the exact function of almost all of these epimerases remains unknown. Two of the currently known algal MC5Es have been functionally described: in *Saccharina japonica,* a recombinant epimerase (SjC5-VI) epimerizes M to G (96). In *Ectocarpus siliculosus*, a mannuronan C5-epimerase MEP13-C5 is thought to epimerize block MM regions, although its exact function is not completely clear (97).

The cell wall architecture in fucoid zygotes has previously been observed by TEM. These studies described that during zygote development, the wall consisted of a single fibrous layer and, as development progresses, several wall layers were observed (86,91,98,99). Between these layers, the spatial arrangement of the fibrils is different, and a recent study has shown that each of these layers seems to have a different alginate composition, which might have different physicochemical properties (100). In addition, this study has shown that removing calcium from the growth medium seems to affect the alginate composition and wall integrity, suggesting that, similarly to our findings, alginate might play a mechanical role in the brown algal cell walls.

### Transcriptome-wide gene expression changes follow known developmental progression

There have been several reports on transcriptomic analyses in brown algae (95,101–109); this study, however, represents the first temporal gene expression analysis of brown algal embryo development. Our transcriptome covers 4 developmental stages during *Fucus* embryogenesis starting just after fertilization and ending at 10DAF. In our gene ontology analysis, several GO terms were enriched in specific time points. In the early development (7h and 1DAF) highly expressed genes are related to cell cycle, cytoskeleton and chromosome segregation. These enrichments align with the developmental processes observed in early *Fucus* embryogenesis: the *Fucus* zygote exhibits spatial distribution of cellular components that are necessary for embryo polarity and the first asymmetrical cell division by activating the cytoskeletal machinery and calcium fluxes (26,34,62,63).

After the initial cell divisions, we show that at 3DAF there is a high expression of genes related to protein metabolism and translation, suggesting that the embryo activates its translational machinery around this stage. This is further supported by a previous study from Galun and Torrey (1969) that demonstrated blocking protein synthesis at 3DAF led to blockage of apical hair production and thallus expansion. Another observation from our transcriptome-wide analysis was the enrichment of photosynthesis-related genes at both 3 and 10DAF. This activation of photosynthesis genes correlates well with physiological and biochemical observations previously reported; zygotes can photosynthesize immediately, but the intensity of photosynthesis increases in several days old embryos (65,66,110–112). At 10DAF, the GO term enrichment is similar to 3DAF indicating that the ‘mature’ gene expression pattern becomes evident very early in fucoid embryo development. Tarakhovskaya et al. (111) have shown that the metabolism of embryos 6-9DAF is similar to adult algae. Since RNA abundance is not always directly correlated with protein levels, it is likely that the initiation of gene expression may occur before changes in metabolism would be evident.

To conclude, our transcriptome analysis has shown that gene expression follows the progression of embryogenesis, with GO terms related to chromosome segregation, cytoskeleton activity and cell cycle being enriched during active cell polarization and first cell division, and protein synthesis, translation and photosynthesis being enriched in later stages, as the embryo starts maturing. We fully expect that there is a wealth of data within the transcriptome that will prove relevant and useful to the community in future studies.

## Materials and methods

### Sample collection and processing

Adult *Fucus serratus* samples were collected in Rottingdean (East Sussex, United Kingdom) during winter months between November 2015 and May 2017. After collection, they were transported in seawater to the Sainsbury Laboratory (Cambridge, UK) and kept at 4°C. Fertile adult samples were rinsed with tap water and processed as follows: each receptacle was first identified as a male or female by checking for antheridia or oogonia, respectively. The receptacles were separated, wrapped into damp tissue paper and aluminum foil (darkness) and kept at 4°C for further use up to 2 weeks.

### Fertilization

The female receptacles were taken out of the 4°C, washed, cut into small segments, placed into beakers with filter-sterilized artificial seawater (ASW, Tropic Marin Sea Salt; Tropic Marin, Germany) and left to release the eggs for approximately 1 hour. The tissue was then removed and the egg mixture was filtered through a 100 µm mesh to eliminate oogonia and leftover pieces of adult tissue. The male receptacles were then taken out of the 4°C, washed, cut into small segments and added to the egg mixture. After 15 minutes, the male segments were removed and the egg/sperm mixture was filtered through a 40 µm mesh to remove the sperm. The fertilized eggs were then placed in droplets of ASW on Multitest 8-well slides (Vector Laboratories, USA) and placed in the incubator. After allowing them to settle for 6 hours, the eggs were flooded with ASW to completely cover the slides and cultured under a unilateral light overnight followed by 12:12 hour day-night cycle, 16°C, 60 µmol m^-2^ s^-1^.

### Light microscopy and measuring length/growth rate

To measure their growth in time, the embryos were cultured under the previously mentioned conditions and imaged using a VHX 5000 microscope (Keyence Ltd, UK) for a proscribed number of consecutive days, depending on the experiment. The images were then processed using ImageJ (113) software where the length of the embryos was measured (drawing a segmented line along the middle of the embryo body from the tip of the rhizoid until the top of the thallus). Growth rate was determined via the difference between sequential daily lengths using the formula R = 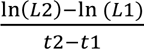; where L2 and L1 are embryo lengths at time t2 and t1 (114).

### Quantifying cell divisions

*Fucus* embryos were cultured on slides and one slide was taken daily for staining and confocal imaging. The embryos were stained with Calcofluor White (18909, Sigma-Aldrich, USA) for 5 minutes, rinsed thoroughly with ASW and imaged under a Leica SP8 confocal microscope (Leica Microsystems, Germany; ex = 370 nm, em= 420 nm). Confocal images were then processed using MorphoGraphX software; www.MorphoGraphX.org; (53)) to extract the information about individual cell surface areas per sample. Briefly, the z-stack output from the confocal microscope was loaded into the MorphoGraphX software as a .tif. The images were first blurred by averaging, after which a global shape of the object was created. Following this, the surface was extracted from this shape as a mesh formed of triangles which were then subdivided and smoothed. The confocal fluorescence signal was then projected onto the mesh after which individual cells were seeded and segmented. Surface areas of each of the cells were analyzed and surface area heat maps were created for individual embryos. Further analysis was conducted using the statistical software R (http://www.R-project.org/").

### Identifying tip growth with Calcofluor White staining

To identify the incorporation of new material into the embryo tip, embryos were stained with Calcofluor White (18909, Sigma-Aldrich, USA) for 5 minutes, rinsed and imaged under a Leica SP8 confocal microscope. The same embryo was imaged again for two consecutive days to locate the incorporation of new wall material.

### Atomic force microscopy (AFM)

Embryos were fertilized, cultured and grown as described above on glass slides and used when reaching the stage of interest: 1DAF, 3DAF or 10DAF. They were covered with a droplet of water and placed under the atomic force microscope. The AFM data were collected using a NanoWizard AFM with a CellHesion (JPK Instruments AG, Germany). The measurement of wall properties was done placing the embryos in ASW. A 0.5 N/m stiffness cantilever with a 10 nm pyramidal tip (Nanosensors, PPP-CONT, Windsor Scientific Ltd., UK) was used with an applied force of 150nN (setpoint). The stiffness of all samples was determined by indenting with the tip over the whole embryo in 100 µm × 100 µm squares with the indentation depth of between 1 and 3 µm. Each force-indentation curve was processed using the JPK Data Processing software (JPK Instruments AG, Germany) to determine the stiffness per indentation point. Stiffness was presented as a heat map; areas of interest were extracted for quantitative analysis by picking points in a line along the middle of the embryo, from the tip of the rhizoid to the top of the thallus, using a custom MatLab-based script (available upon request).

### Alginate immunolocalization

Embryos were fertilized and cultured as above on multi-test 8-well slides (Vector Laboratories, USA) and taken when reaching the stage of interest (1, 3, and 10DAF). They were fixed overnight in ASW containing 2% formaldehyde and 2.5% glutaraldehyde and washed 3 times for 15 minutes with ASW, followed by a rinse in phosphate buffered saline (PBS; 2.7 mM KCl, 6.1 mM Na_2_HPO_4_, and 3.5 mM KH_2_PO_4_). The samples were incubated in a blocking solution of 5% milk for 2 hours. They were then rinsed with phosphate buffered saline and incubated in the 60 µl of 1/5 (in 5% milk) monoclonal primary antibody for 1.5 hours. After the incubation, the slides were washed with PBS 3 times for 5 minutes each, followed by the incubation in the 60 µl of 1/100 (in 5% milk) IgG-FITC secondary antibody (F1763, Sigma-Aldrich). This was followed with a 5×5 minute wash in PBS, after which the samples were mounted in Citifluor (Agar Scientific, UK), covered with a coverslip, sealed and imaged under a Leica SP8 confocal microscope (Leica Microsystems, Germany; ex=490nm, em=525nm)

### RNA extraction, cDNA synthesis, and RNA sequencing

Total RNA of 3 biological replicates of embryos from 7 hours (H), 1 day (D), 3 days (D) and 10 days after fertilization (DAF) was extracted using the PureLink Plant RNA Reagent following manufacturer’s instructions. The integrity of RNA samples was checked by Agilent 2100 Bioanalyzer (Agilent Technologies, USA) and the quantity was assessed using NanoDrop 1000 spectrophotometer (Thermo Fisher Scientific, USA) and Qubit 2.0 Fluorometer with RNA High Sensitivity assay (Thermo Fisher Scientific). cDNA libraries were generated using TruSeq LT DNA Sample Prep Kit (Illumina, USA) according to the manufacturer’s instructions with the following modifications: the beads used were home-made SeraPure beads (115) instead of AMPure XP beads. The library sequencing was performed on a NextSeq 500 using paired-end sequencing (2×76 cycles) with NextSeq 500/550 High Output v2 kit (Illumina, USA).

### *De novo* transcriptome assembly and GO term analysis

A total of 65-160 million paired-end reads (75×75bp) were generated for each of the 12 sequenced libraries (14 samples in total; sequencing was done on two separate flow cells and to remove possible sequencing bias, two libraries were sequenced in both runs). To reconstruct *F. serratus* transcriptome, samples were pooled together from all four time-points (7H, 1D, 3D, and 10DAF). Initial read quality assessment was done with FastQC (Babraham Bioinformatics, www.bioinformatics.babraham.ac.uk/projects/fastqc/). Adaptors were removed using CutAdapt (116). Reads were further subjected to quality control using Trimmomatic (minimum read length = 60). The quality parameters for the library were assessed using FastQC. The resulting filtered reads were subjected to *de novo* assembly with Trinity (trinity v2.4.0) on a high-RAM server with minimal k-mer coverage = 2 and k-mer length = 25. *In silico* read normalization was used due to a large number of input reads, in order to improve assembly efficiency and to reduce run times (117). Trinity analysis resulted in 127,489 transcripts, accounting for 70,824 nominal genes with an average length of 780bp. It was suspected that the Trinity assembly yielded a significant amount of duplication beyond the isoform level (genes are called as nominal even with a very high sequence similarity) and that many genes were fragmented. To improve the assembly and overcome the fragmentation issue, Salmon and CD-HIT were used to collapse the Trinity genes into larger unigene groups (118, 119). Analysis of the percentage of unigene collapse with mappability started severely decreasing when the genes were collapsed for more than 80% of nucleotide identity (Fig. S4). The level of 80% was then chosen as the new database for unique genes. Salmon filtering method resulted in 42,176 contigs, with the length ranges of 201 to 17,616 nt, and a mean length of 1,163 nt. It was predicted that 24,691 genes had ORFs, and 67% (16, 597) shared sequence homology with a representative of the NCBI database. Gene abundances were analyzed following the general outline of R Bioconductor package Sleuth (120). All DEGs were then used for GO term analysis in the Gene Ontology database (http://geneontology.org/). To remove redundancy in the number of similar GO terms and choose a representative subset of the terms we used the REVIGO algorithm with allowed similarity of 0.7 and the default SimRel semantic similarity measure (121). GO terms with p-values ≤0.001 were defined as significantly enriched.

## Acknowledgments

The authors thank Dr. Anna Gogleva and Dr. Sebastian Schornack for their help with the initial *Fucus* embryo transcriptome assembly and Dr. Thomas Torode for help learning brown algal cell wall immunolocalizations. We also thank Dr. Paul Knox and Dr. Cécile Hervé for the gift of BAM antibodies, which are now available at Sea Probes (http://www.sb-roscoff.fr/en/seaprobes). The Braybrook group in Cambridge (UK) was funded by The Gatsby Charitable Foundation (GAT3396/PR4, S.A.B) and the Ralph Lewin – F. E. Fritsch Prize Studentship (M.L). The Braybrook group at UCLA is funded by The Department of Cell, Molecular and Developmental Biology and The College of Life Sciences (S.A.B); At UCLA this work was majorly supported by the U.S. Department of Energy Office of Science, Office of Biological and Environmental Research program under Award Number DE-FC02-02ER63421 and the US Department of Energy (Biological and Environmental Research (BER), the Biological Systems Science Division (BSSD); M.L, S.A.B, M.P).

## Supplementary data

**Supplemental Figure 1.**
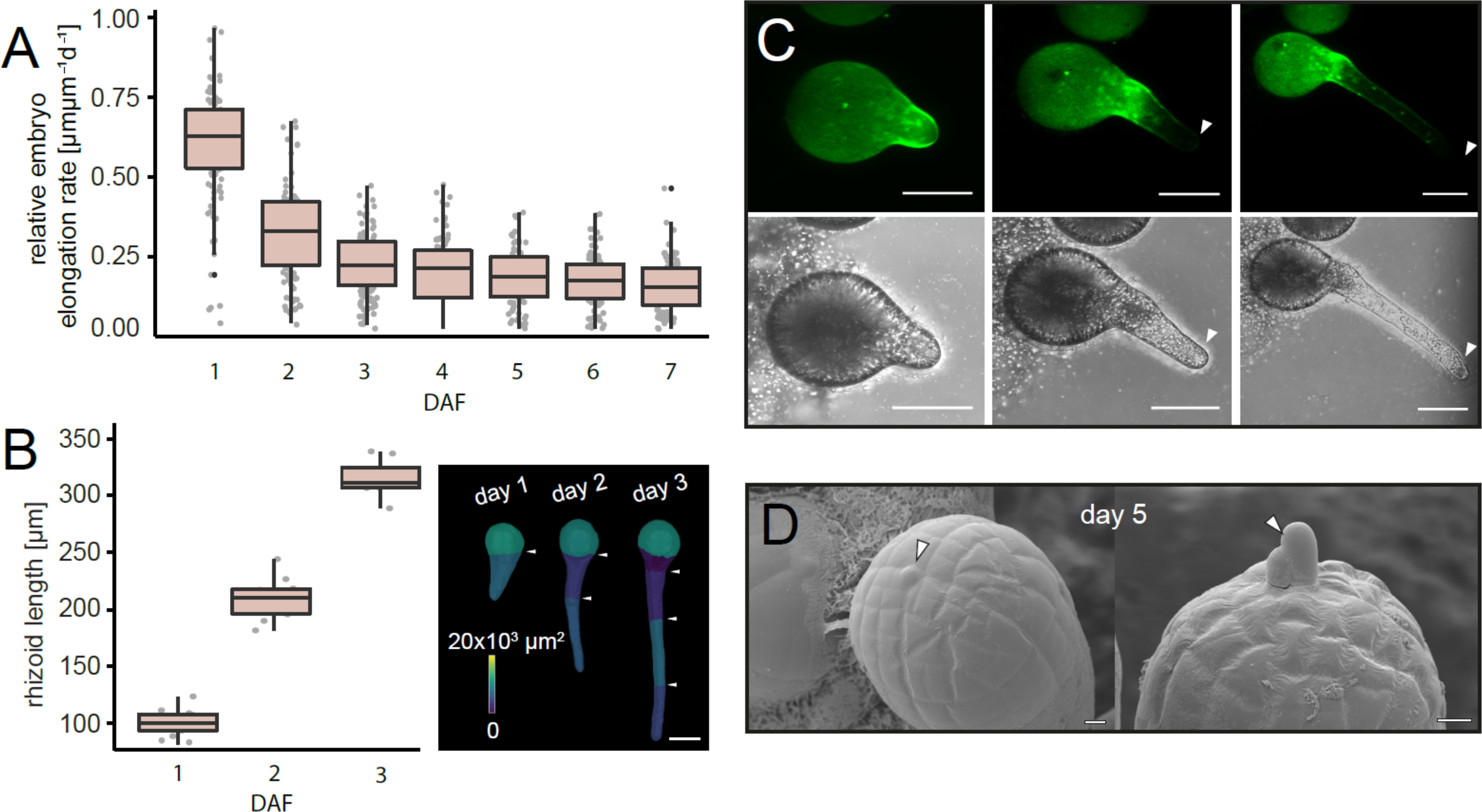
Rhizoid and thallus growth in the *F. serratus* embryo. (A) Relative growth rate decreases during the first 7 days of development and becomes constant around day 5. (B) Rhizoid length increases significantly during the first days of embryogenesis; few cell divisions can be observed. Scale bar 50 µm. (C) Calcofluor White staining of the embryo cell wall at 24h after fertilization (AF) after which the stain was removed from the medium. Embryos were imaged 48 and 72h AF. Images represent an embryo with the retained stain and the new part of the unstained wall at the tip (white arrowhead). Bright-field images of the three stages. Scale bar 50 µm. (D) Initiation of apical hairs around day 5 (white arrowheads) indicates the start of active meristematic growth in the embryo thallus. Scale bar 10 µm.

**Supplemental Figure 2.**
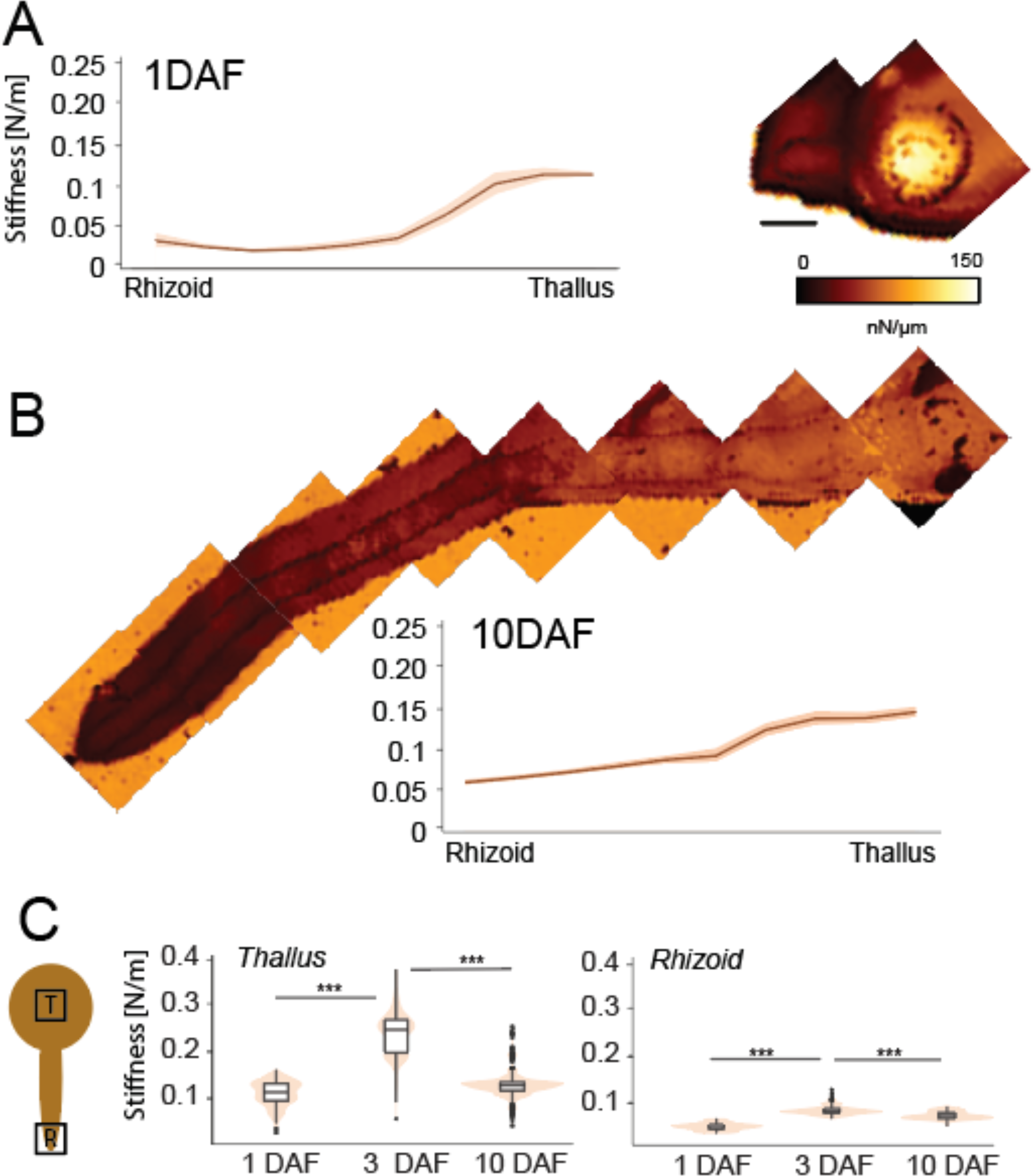
Atomic Force Microscopy analysis for 1 and 10 DAF. (A) Stiffness map and corresponding graph for 1DAF (B) Stiffness map and corresponding graph for 10DAF, both indicating rhizoid area as less stiff than the thallus. Scale bar 50 µm. (C) Stiffness differences in thallus (A) and rhizoid (B) during embryo development. Thallus cell exhibits the highest stiffness during early cell division stage (3DAF), but decreases after cell expansion takes place (10DAF). In the rhizoid, the stiffness increases in the early development but becomes reduced at later stages.

**Supplemental Figure 3.**
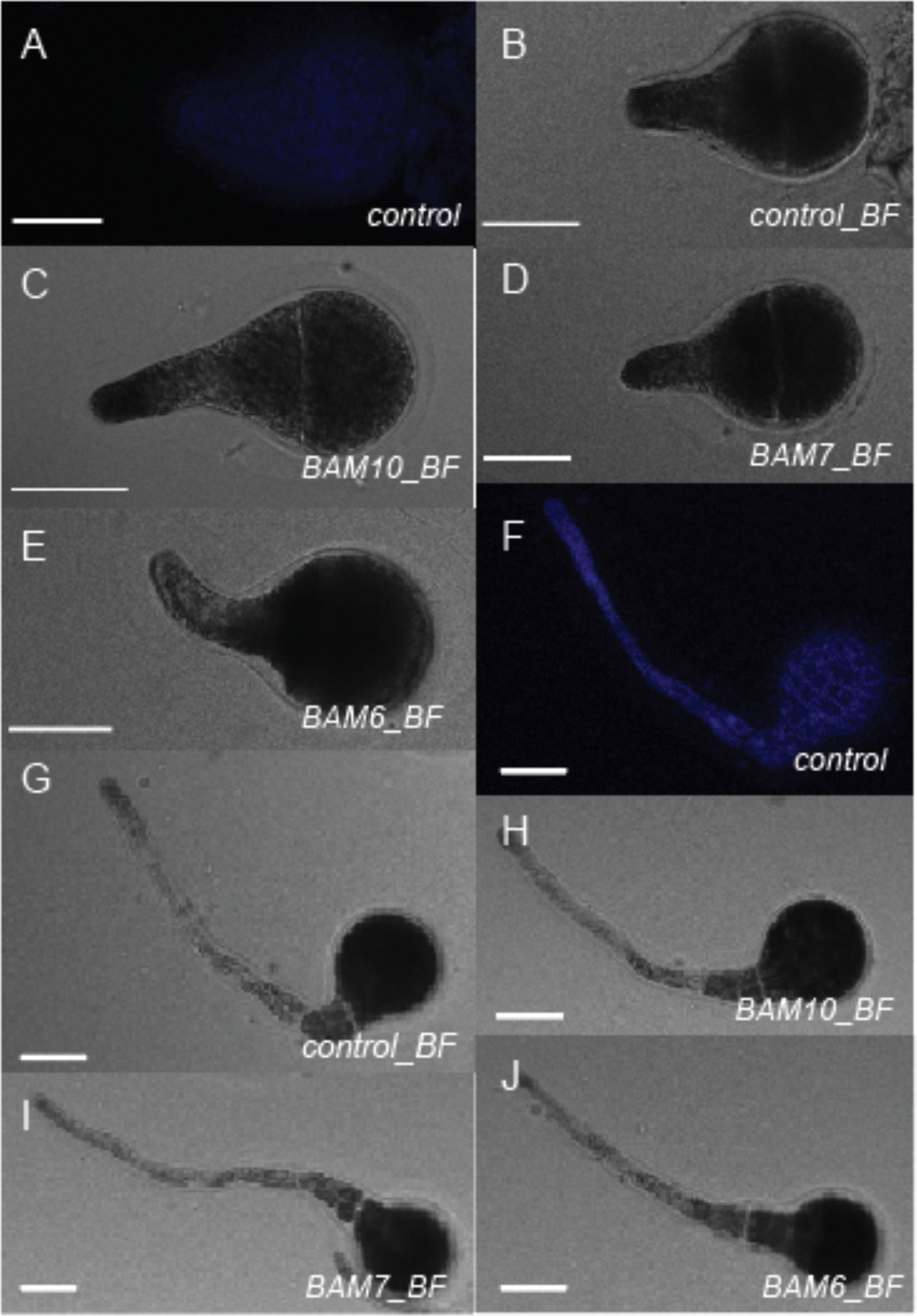
Control images for alginate immunolocalization. (A) No primary antibody 24h AF control, (B-E) Bright-field images of 24h old embryos, (F) No primary antibody 72h AF control, (G-J) Bright-field images of 72h old embryos. Scale bar 50 µm.

**Supplemental Figure 4.**
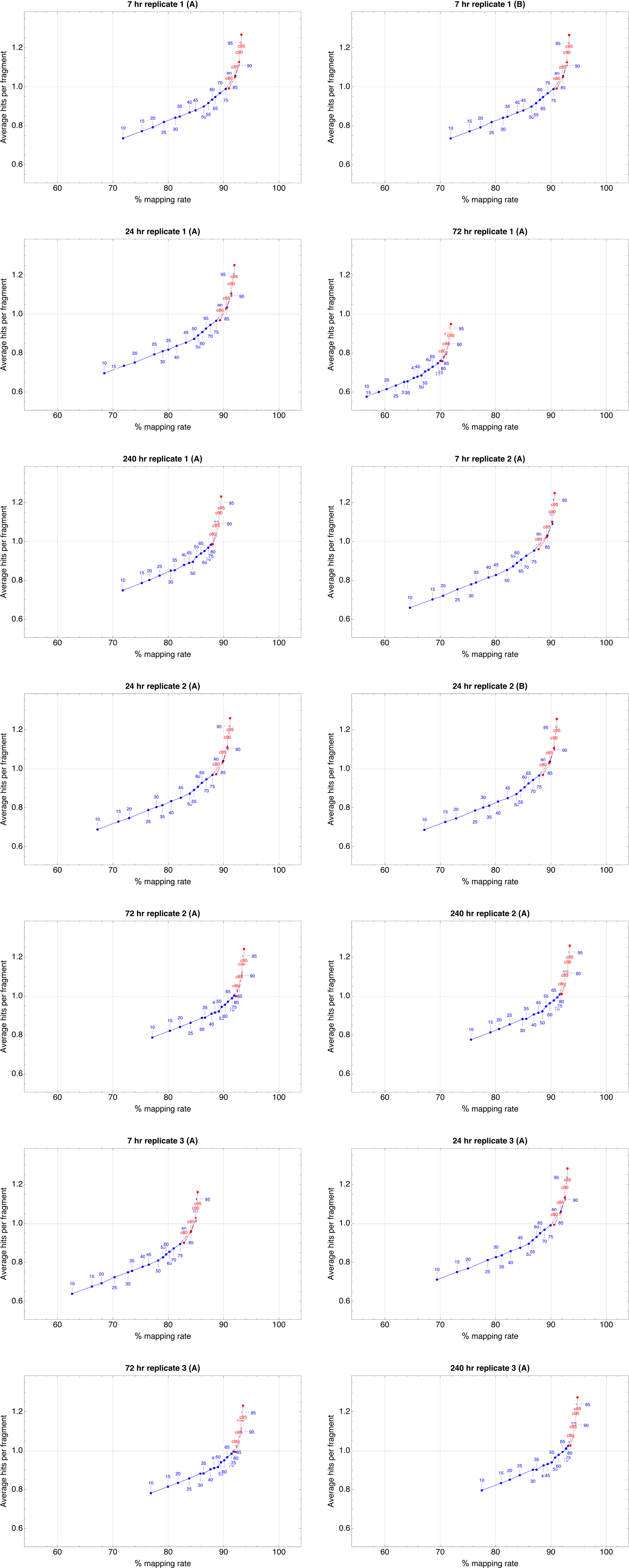
CD-HIT clustering reads and the corresponding mapping rate. CD-HIT (red) and PSI-CD-HIT (blue). *SEE END OF DOCUMENT*.

**Supplemental Figure 5.**
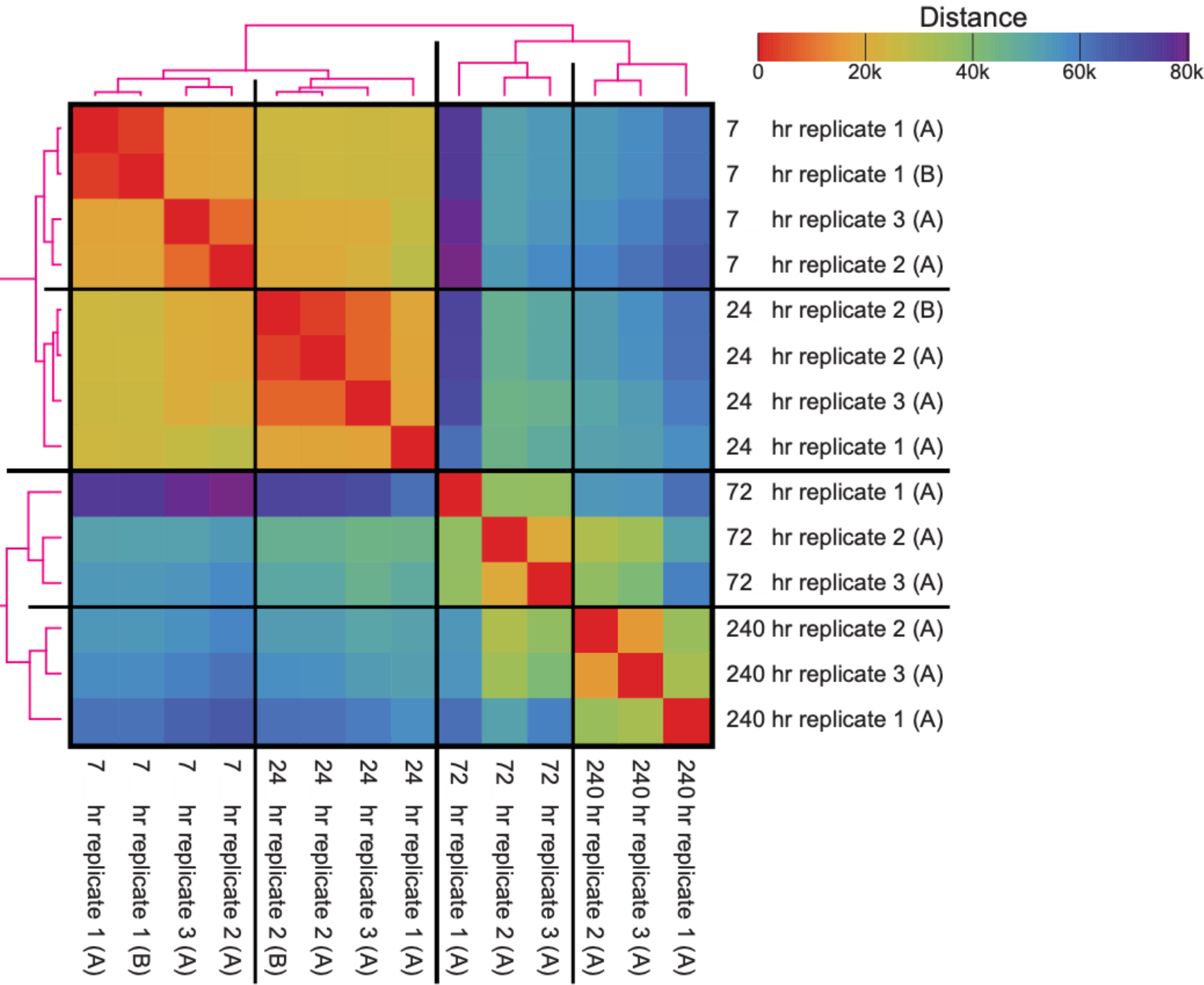
Salmon normalized expected reads (14 samples by 72,788 transcripts; two libraries were done in duplicate to assess sequencing bias, see Materials and Methods), reduced to 27,110 transcripts that have at least one sample with >= 200 normalized counts. Transformed by log2(*+0.5), hierarchically clustered by Manhattan distance with complete linkage.

**Supplemental Figure 6.**
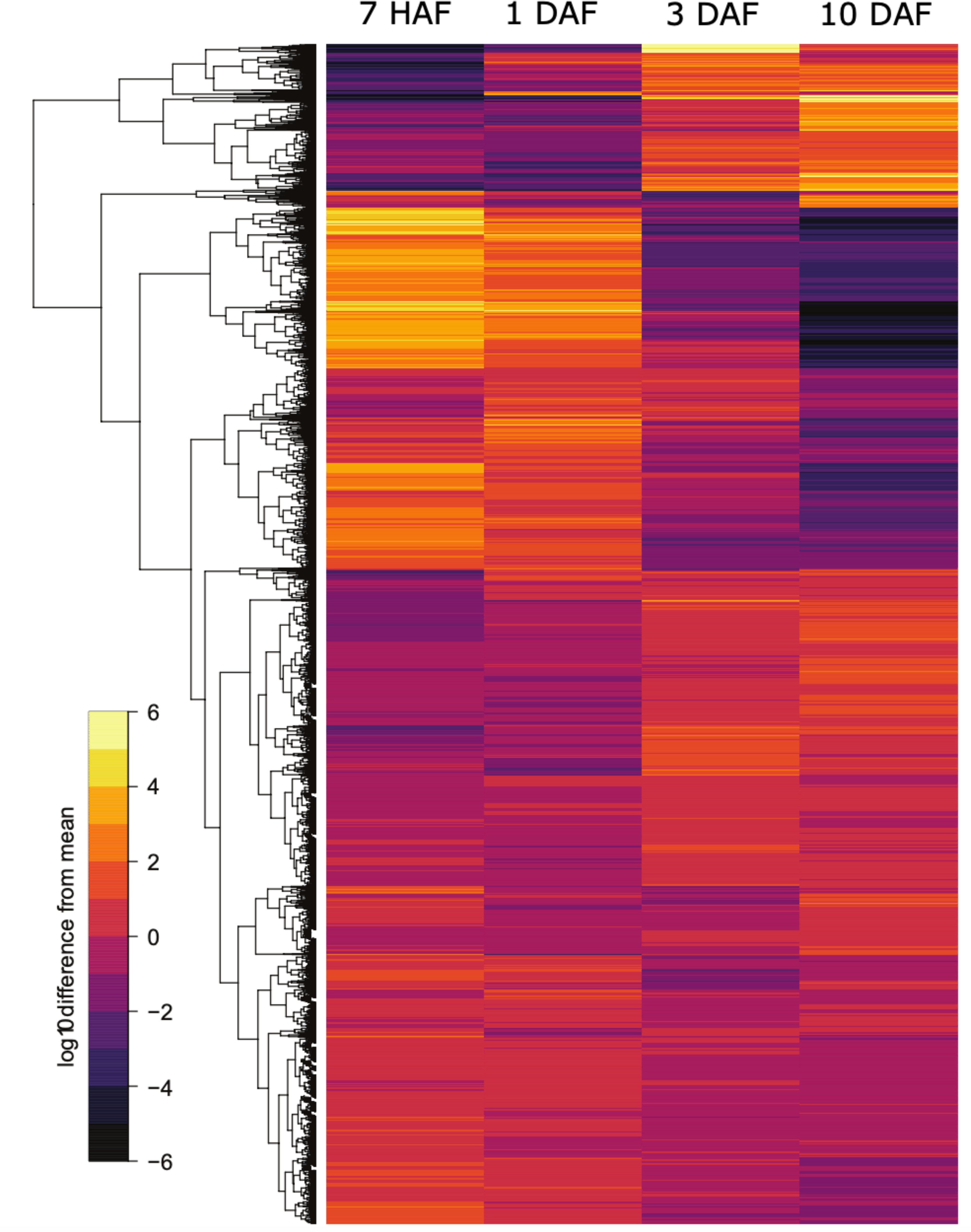
Gene expression during *Fucus* embryo development; depicting all genes differentially expressed in at least one time-point (7HAF, 1DAF, 3DAF, and 10DAF; p-value ≤ 0.0001).

**Supplemental Figure 7.**
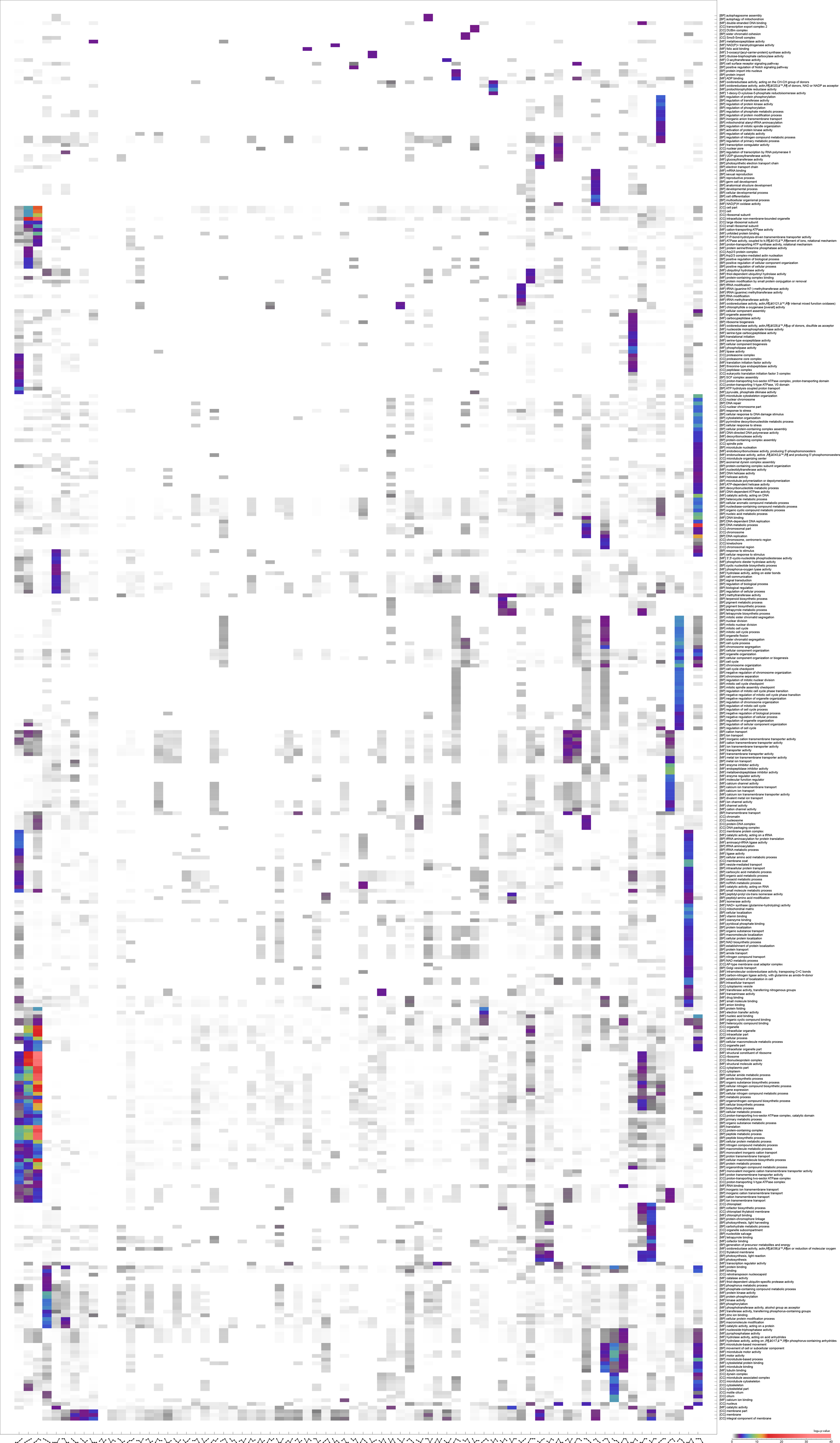
Heat map of Gene Ontology enrichments; full list of all rows and columns containing *p*-value ≤ 0.0001. *SEE END OF DOCUMENT*

**Supplemental Figure 8.**
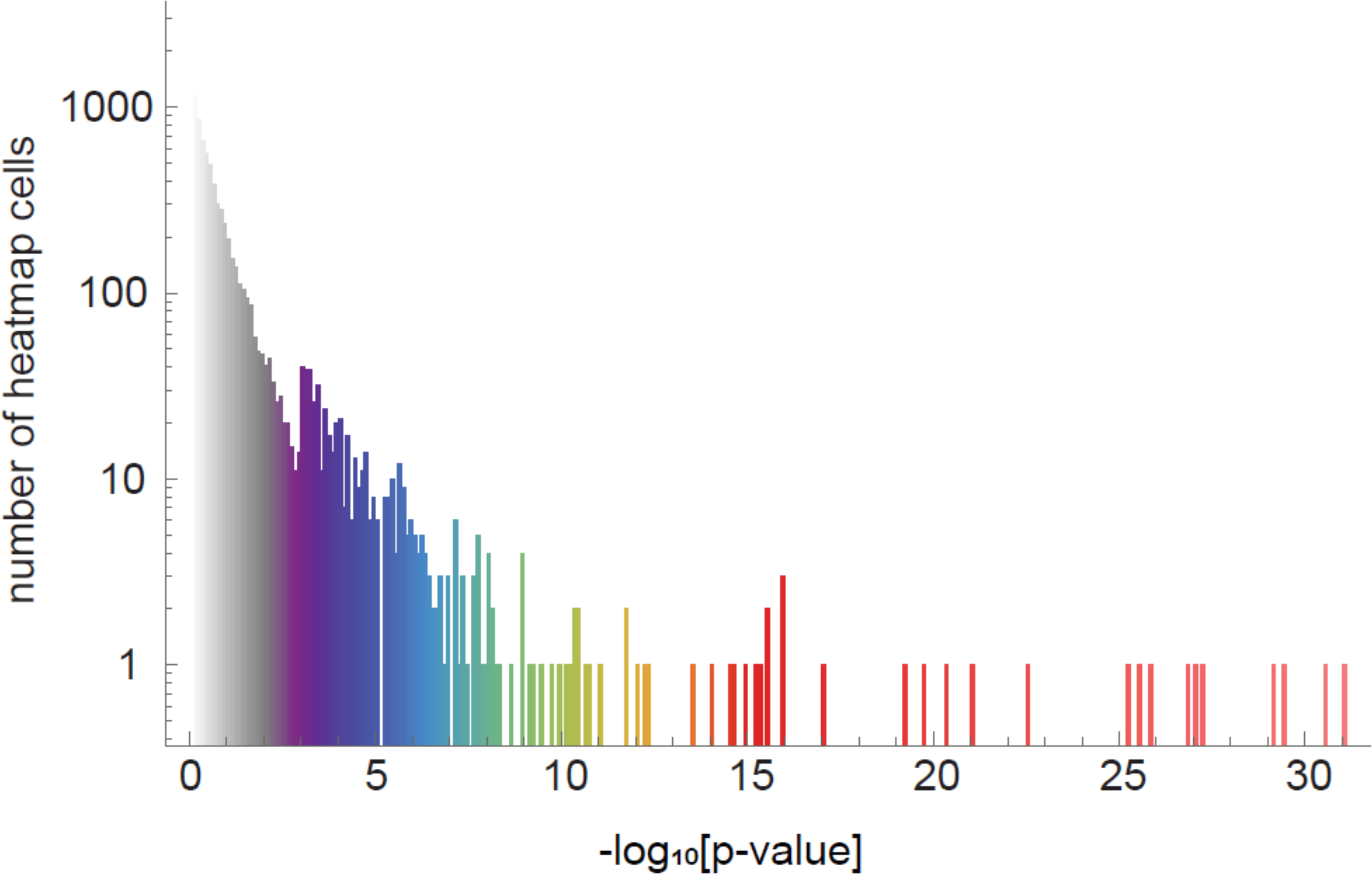
Heat map representing GO terms across time classes and their enrichment significance. There are 74 (time classes) by 381 (GO terms) = 28,194 cells in the heat map and the histogram represents a single value shown in each cell (-log_10_ enrichment p-value). The x-axis shows bins for the -log_10_[p-value], the y-axis shows the number of heat map cells in the bins. Purple to red color refers to the 381 GO terms that are cut down from the 2,339 GO terms enrichment was computed on; p-value <= 0.001 in at least one of the 74 time-classes. White and gray colors mark the GO terms with p-value > 0.001.

**Supplemental Table 1. Cell wall biosynthesis genes and their expression during *Fucus* embryogenesis.** List of Trinity genes with best NCBI hit or locally performed BLAST with *Ectocarpus* wall biosynthesis genes against a local *Fucus* transcriptome database. Gene expression levels presented as the deviation from the mean of all time points with the log_10_(q-value).

